# Infant gut bacteriophage strain persistence during the first three years of life

**DOI:** 10.1101/2023.08.07.552389

**Authors:** Yue Clare Lou, LinXing Chen, Adair L. Borges, Jacob West-Roberts, Brian A. Firek, Michael J. Morowitz, Jillian F. Banfield

## Abstract

Bacteriophages are key components of gut microbiomes, yet the phage colonization process in the infant gut remains uncertain. Here, we established a large phage sequence database and used strain-resolved analyses to investigate phage succession in infants throughout the first three years of life. Analysis of 819 fecal metagenomes collected from 28 full-term and 24 preterm infants and their mothers revealed that early-life phageome richness increased over time and reached adult-like complexity by age three. Approximately 9% of early phage colonizers, mostly maternally transmitted and infecting *Bacteroides*, persisted for three years and were more prevalent in full-term than in preterm infants. Although rare, phages with stop codon reassignment were more likely to persist than non-recoded phages and generally displayed an increase in in-frame re-assigned stop codons over three years. Overall, maternal seeding, stop codon reassignment, host CRISPR-Cas locus prevalence, and diverse phage populations contribute to stable viral colonization.

## Introduction

The early-life gut microbiome assembly has a significant impact on infant development and health^1–3^. The infant gut microbiome, with an initially low species diversity and a high turnover rate^4^, undergoes several distinct microbiome states before reaching a compositionally stable and diverse stage by age three^5, 6^. This gut microbiome succession is crucial for infant development, and disruption of this process can increase the likelihood of developing diseases, such as asthma and atopy, later in life^7–9^.

Metagenomics analyses on prospective birth cohorts have revealed detailed insights into early-life succession dynamics of gut bacteriomes^3, 10–14^. However, much less is known about the colonization and persistence of bacteriophages (phages) in infants^15^. Phages are viruses that prey on bacteria for reproduction, which typically results in the lysing of the bacterial host cell^16, 17^. Temperate phages have the option of integrating into the host chromosome, forming a prophage, and replicating with the host rather than killing it^18–20^.

In the adult human gut, it is estimated that phages and bacteria exist in a ∼1:1 ratio, although the actual number can be potentially higher (i.e., in the mucus layer)^21, 22^. The prevalence of phages in the human gut suggests constant and frequent interactions between phages and bacteria. Indeed, phage parasitism has been found to impose a strong force driving the diversification of bacterial populations of the same strain or species across different environments, including the human gut^23, 24^.

The adult gut microbiome is compositionally and functionally stable^25, 26^. Year-long gut colonization is not only seen in self-replicating microorganisms like bacteria but also in phages^27, 28^. At the community level, this contrasts with the developing infant gut microbiome, thus raising the question of the existence of long-term persisting phages in infants. Phages cannot replicate on their own and instead, rely on bacterial hosts for survival^17^. Our earlier work revealed a small percentage of bacterial strains could persist in infants for at least one year^13^. Given the development of viromes likely parallels that of bacteria^29, 30^, it is possible that certain phages could also persist in infants for one year or longer. These long-term persisting phages could possibly play pivotal roles in shaping the gut microbiome assembly. It is thus important to analyze phage strain persistence, as well as factors contributing to their stable colonization, especially given the infant gut microbiome is highly dynamic.

The acquisition of phages occurs shortly after birth^29, 30^. Given the significant selective pressure they impose on bacteria, it is plausible that phages play an indispensable role in shaping the assembly trajectory of the infant gut microbial community. Early-life virome metagenomic studies to date not only expanded the known viral species^31^ but also offered an overview of the viral colonization and assembly during infancy^29, 30, 32, 33^. However, these studies focused on viral assembly at the species-and/or genus-level, which typically consists of genomically different phage strains with potentially distinct functional capacities and/or host targets^34–36^. Longitudinal bacterial studies have revealed strain fluctuations within individuals over time, despite showing stable compositions at a higher taxonomic level (i.e., genus)^3, 13, 37^. Similarly, a handful of virome studies have shown genomic evidence of within-person evolution of the same gut phage population^27, 34, 36–38^. In some cases, a single point mutation in the tail fiber protein can enable phages to switch hosts^34^. It is thus important to examine the succession dynamics of infant viromes at a finer resolution.

Here, we investigated early-life gut phageome assembly dynamics using genome-resolved metagenomics. We recovered over 30 thousand medium-to high-quality phage contigs using 819 fecal samples from preterm and full-term infants, collected from birth to age three, and from their mothers at birth and after three years. Over 40% of the assembled phage sequences did not share any close relatives with those in the published human gut phage databases, supporting the importance of having project-specific phage databases. We subsequently applied rigorous strain-resolved analyses to investigate phage colonization dynamics during infants’ first three years of life and phage transmission between mothers and infants. We determined that maternal origin, bacterial host persistence, genetic code expansion, and high population diversity all contributed to phage persistence in infants.

## Results

### *De novo* construction of human gut phage database

In this study, we followed 28 full-term and 24 preterm infants, along with their mothers, from birth to up to three years of age (Figure S1 and Table S1). We grouped the infant fecal samples into six time windows based on the infants’ chronological ages at the time of collection (Table 1 and Figure S1). A median of 11 fecal samples was collected from each infant (Figure S1 and Table S2). Up to two maternal fecal samples were collected, with one being around birth and one when the infant turned three years old (Figure S1). In total, 735 and 84 fecal DNA samples from 52 infants and 42 mothers, respectively, were extracted and subjected to deep metagenomic sequencing (∼8.15 tera base pairs (Tbp) of total sequence data in the form of 150 bp paired-end reads).

**Table 1.** Three-year sampling time windows.

| Time windows | Months | Num. Individuals | Num. DNA samples |
| --- | --- | --- | --- |
| Mom1 | ~Birth | 39 | 53 |
| Mom2 | ~36 months post birth | 31 | 31 |
| W1 | Months 0-2 | 52 | 291 |
| W2 | Months 3-4 | 50 | 110 |
| W3 | Months 8-12 | 52 | 129 |
| W4 | Months 15-16 | 38 | 39 |
| W5 | Months 20-25 | 45 | 87 |
| W6 | Months 30+ | 40 | 79 |

Reads were *de novo* assembled, and a total of 32,401 bacteriophage contigs and 28,448 microbial draft genome bins were recovered. Subsequent genome dereplication at 98% whole-genome average nucleotide identity (gANI) yielded 8,424 phages and 1,951 microbial genomes, which represent unique “subspecies” (Methods)^13^. The dereplicated phage genomes had an average genome length of 45.8 ± 25.5 kb, with the minimum and maximum, both circular, being 4.6 kb and 394 kb, respectively.

Detection of identical strains was achieved using inStrain^39^ (Methods). We applied the same population-level ANI (popANI) cutoff, 99.999%, as our previous study, when identifying near-identical bacterial strains^13, 39^. Phages have an approximately 1000x faster mutation rate than bacteria^40^, we thus lower the popANI threshold to 99% to allow the identification of within-host *de novo* mutated phage strains and/or recent phage-transmission events (e.g., vertical transmission from the mother to the infant) (Methods). Data contamination was assessed as previously described^41^, and contaminated samples, including all samples collected from one preterm infant (infant ID “#60”), were removed before proceeding with the data analysis (Table S2).

### Expanding known human gut phage species

Recent publications of five large-scale human gut phage studies have generated large public phage sequences databases^31, 42–45^. To examine the novelty of our reconstructed phage genomes, we clustered our 8,424 dereplicated phage representative sequences and 285,223 sequences from these five phage databases at 95% ANI over 85% of the length (Methods). Our 8,424 dereplicated phage genomes clustered into 7,398 phage species. Notably, 3,343 phage species from our study (∼45.2%), corresponding to 3,397 phage genomes, did not cluster with any sequences from these reference phage genome sets, suggesting the novelty of our constructed phage genomes. This result underlines the importance of assembling person-specific phage sequences, rather than mapping reads or contigs to public reference databases, for the characterization of virome development.

### Early-life gut phage community overview

For both pre-and full-term infants, their gut phage alpha diversity increased as infants matured (Figure 1A) (Spearman correlation coefficient r = 0.69 for Shannon diversity index; p = 1.23e-29) (Methods). A similar trend was seen when measuring the phage alpha diversity analysis using Phanta, an assembly-free, k-mer-based phage identification tool^46^ (Figure S2) (Spearman correlation coefficient r = 0.77 for phage richness normalized by sequencing depth; p = 1.34e-55). Notably, the increase in phage alpha diversity over time positively correlated with that of bacteria (Figure 1B) (Spearman correlation coefficient r = 0.77 for Shannon diversity index; p = 1.89e-55).

**Figure 1.**
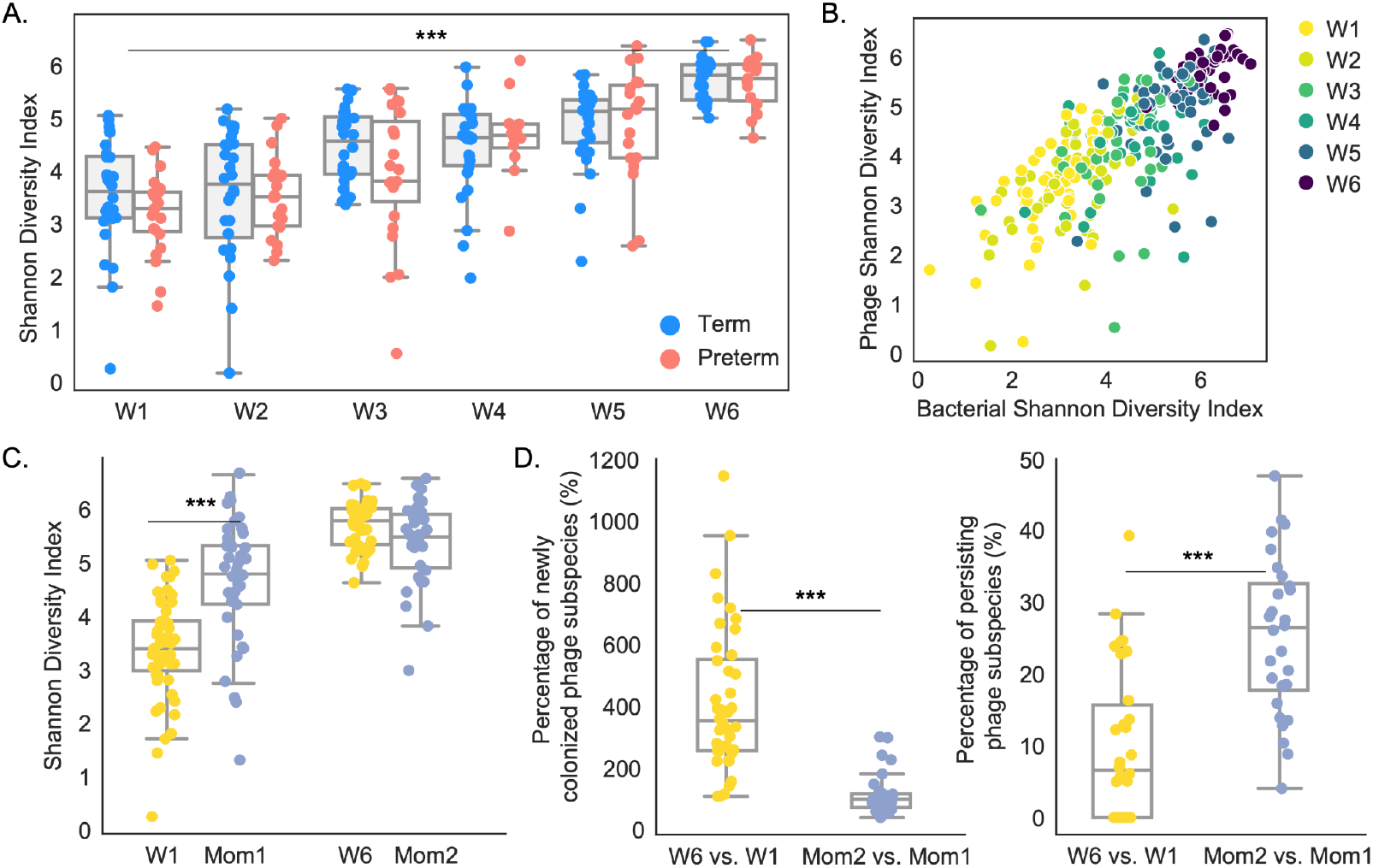
The infant phageome complexity was comparable to mothers by age three. (A) Gut phage alpha diversity, measured via the Shannon Diversity Index, in full-term and preterm infants over time. Full-term and preterm infants are shown in sky-blue and salmon-red, respectively (*** = p < 0.001). (B) Gut bacterial (x-axis) versus phage (y-axis) alpha diversity, measured via the Shannon Diversity Index, in infants from birth to age three. Each dot represents an infant and is colored by age. (C) Gut phageome complexity comparison between infants and mothers when infants were less than 2 months old (W1 vs. Mom1) and when infants were 3 years old (W6 vs. Mom2). The phageomes of infants and mothers are shown in gold and cornflower blue, respectively (*** = p < 0.001). (D) Percentages of newly colonized (left panel) and persisting (right panel) phage subspecies detected in the final sampling window (W6 or Mom2) when compared to the initial window (W1 or Mom1). The color scheme is the same as panel C (*** = p < 0.001).

Similar to infants, maternal samples collected when infants were age three also had a higher phage alpha diversity than those collected around birth (p = 0.0029; Wilcoxon rank-sum test). This may reflect a temporary reduction in phage diversity due to the process of pregnancy and/or giving birth. Notably, while mothers initially exhibited a higher phage alpha diversity than infants (i.e., W1 vs. Mom1, p = 8.31e-07; Wilcoxon rank-sum test), by age three, the infant phageome diversity reached the same level as their mothers’ (Figure 1C) (W6 vs. Mom2, p = 0.13; Wilcoxon rank-sum test). This observation led us to hypothesize that infants likely had a faster phage acquisition rate than their mothers. Indeed, when examining the final time point (W6 or Mom2), we detected a significantly higher percentage of newly colonized phage subspecies, as well as a significantly lower percentage of persisting phage subspecies, in infants than in mothers, when compared to the initial sampling window (W1 or Mom1) (Figure 1D) (p = 1.44e-10 and 4.29e-6, respectively; Wilcoxon rank-sum test) (Methods).

### Approximately 9% of phages persisted throughout the first three years of life

The identification of persisting phage subspecies in infants and mothers prompted us to search for phage strains that colonized infants for three years. To do so, phages were first classified as early colonizers if they appeared in the infant gut by W1. Using the 99% popANI strain identity cutoff, we further divided these early phage colonizers into “persisters” and “non-persisters” based on their presence (persisters) or absence (non-persisters) at W6 (Methods). This analysis only included the 40 infants who completed the full three-year fecal sample collection period (25 full terms and 15 preterms). In mothers, if a phage was detected in both maternal samples (one around birth and one when infants turned three years old), it was defined as a persister. Otherwise, it was a non-persister. In total, 28 mothers, from whom we collected two 3-year-apart fecal samples, were included in this analysis.

Among the 1801 early-colonizing phages identified in infants, a total of 155 (∼8.6%) persisted for 3 years (Figure S3). These 155 phage strains were found in half of the infants (17 full-term and 3 preterm infants). Persisting phages were detected in all mothers. In total, 711 maternal phage strains (∼18% of the initial colonizers) persisted throughout the 3-year sampling collection period. We found maternal gut phageome to be a critical source of phage persisters in infants, as vertically transmitted phages were significantly more likely to persist in the infant gut than those that were not maternally transmitted (p = 9.90e-95; Fisher’s exact test) (Figure 2A left panel). Interestingly, maternal phage persisters were also more likely to be vertically transmitted to infants than non-persisting maternal phages (p = 4.37e-62; Fisher’s exact test) (Figure 2A right panel). We thus hypothesized that the persistence of phages in both infants and mothers could, in part, be attributed to recurrent re-seeding from both parties.

**Figure 2.**
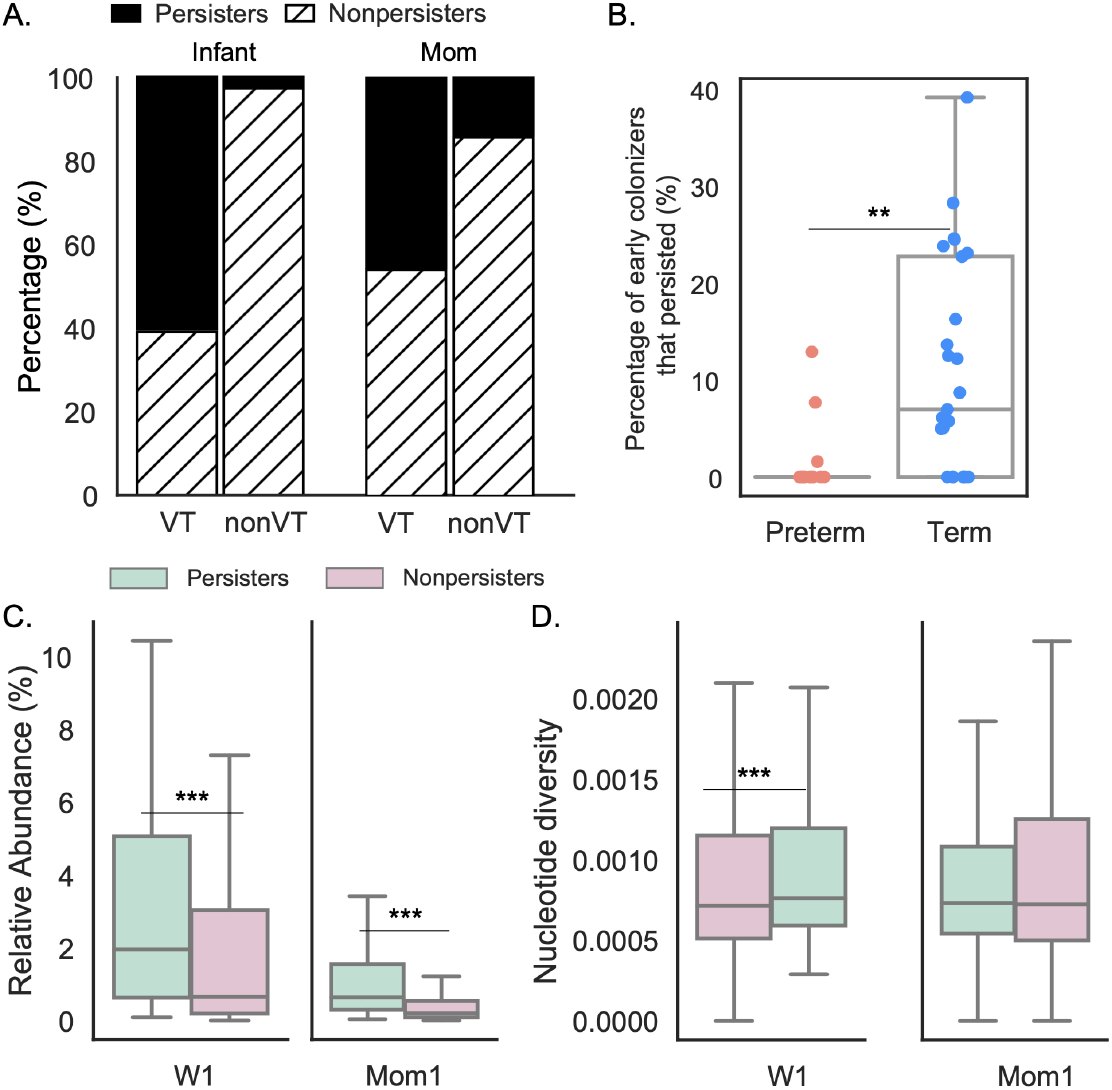
Origin, prematurity, initial colonization abundance, and population diversity influence phage persistence in the gut. (A) Percentages of persisters (solid black) and non-persisters (dashed) in infants (left panel) and mothers (right panel) that were vertically transmitted from the mother to the infant. “VT” stands for vertically transmitted and “nonVT” stands for non-vertically transmitted. (B) Percentage of early colonizers that persisted in the gut microbiomes of preterm and full-term infants. Each dot represents an infant. Salmon-red and sky-blue circles represent preterm and full-term infants, respectively. The box plot shows the interquartile range (IQR) of the percentage of early colonizers that persisted in infants, with the central line representing the median; the whiskers extend from the lower and upper quartiles to 1.5 times the IQR (** = p < 0.01). (C) Relative abundances of persisters (light green) and non-persisters (pink) during the initial sampling window in infants (left panel) and mothers (right panel) (*** = p < 0.001). (D) Nucleotide diversity of persisters (light green) and non-persisters (pink) during the initial sampling window in infants (left panel) and mothers (right panel) (*** = p < 0.001).

Full-term infants were more likely to have phage persisters than preterm infants, regardless of the delivery mode or feeding type (exclusively breastfed or not) (p = 0.0079; Fisher’s exact test). A higher percentage of early colonizing phages also persisted in full-term infants than did so in preterm infants (p = 0.0042; Wilcoxon rank-sum test) (Figure 2B). The size or diversity of the initial phage populations did not confound the comparison of full-term versus preterm infants, as no significant difference was observed in either the total number or the alpha diversity of phage early colonizers (p = 0.28 and 0.14, respectively; Wilcoxon rank-sum test).

At the initial sampling time points (W1 and Mom1), phage persisters in infants and mothers had a significantly higher relative abundance than non-persisters (p = 1.58e-09 and 1.61e-83, respectively; Wilcoxon rank-sum test) (Figure 2C). In infants, we also found phage persisters having a higher nucleotide diversity than non-persisters (p = 1.11e-16; Wilcoxon rank-sum test) (Figure 2D). However, no difference was detected between maternal persisters and non-persisters (p = 0.96; Wilcoxon rank-sum test) (Figure 2D).

Phage persisters in infants were enriched with temperate phages (p = 0.027; Fisher’s exact test) (Methods), suggesting their persistence may be partly due to their ability to stably co-exist with their microbial hosts as prophages. Phage lifestyle difference was not detected between maternal persisters and non-persisters (p = 0.53; Wilcoxon rank-sum test). Nine bacterial genera, with *Bacteroides*, *Parabacteroides*, and *Bifidobacterium* being the three best represented, were more likely to harbor persisting phages than other genera (q < 0.05; one-sided binomial test) (Figure S3 and Table S3) (Methods). Notably, *Bacteroides* and *Parabacteroides* strains themselves were enriched with persisters (q < 0.01; one-sided binomial test) (Table S4) (Methods). Many *Collinsella*, *Megasphaera*, and *Phocaeicola* strains were also persisters (q < 0.01; one-sided binomial test) but were less likely than *Bacteroides* and *Parabacteroides* to have persisting phages (Figure S3; Tables S3 and S4).

### Co-occurrence of phages and their bacterial hosts in infants

The persistence of bacterial strains and phages with predicted hosts from these genera suggests the direct linkage between phage and bacterial persistence. To better understand the relationships between persisting bacteria and phages, we examined the abundance of persisting phages and their predicted bacterial hosts over three years (Methods). Of most interest were phages that persisted without host integration at one or more time points, as these examples are most likely to reveal gut phage-bacteria population dynamics.

We focused on 18 examples for which we could confidently assign bacterial host types using CRISPR spacer matches (Figure S4). Specifically, we allowed host identification if the phage was targeted by CRISPR spacers from a different individual from the same infant cohort or from public databases, so long as the expected host species co-occurred with the phage. In two cases, we could assign a specific bacterial host based on a CRISPR spacer from the genome of a bacterium that coexists with the phage. Both phages are predicted to be temperate. One of these was a *Bacteroides vulgatus* phage from preterm infant #57 (“I57_Bv_phage”), and the other was a *Megamonas funiformis* phage from full-term infant #123 (“I123_Mf_phage”).

I57_Bv_phage was only detected during W1 and W6, while its bacterial host, *B. vulgatus* (“I57_Bv_host”), was detected across all six sampling windows (Figure 3A). Both I57_Bv_phage and I57_Bv_host were maternally transmitted. Read mapping revealed I57_Bv_host CRISPR locus heterogeneity within the population and over time (Figure 3A). Specifically, during W1 and W6, in which both I57_Bv_host and I57_Bv_phage were detected, only a minority of I57_Bv_host populations encoded the CRISPR array, as shown by read mapping across the CRISPR region (Figure 3A). When I57_Bv_phage was not detected (W2-W5), the same CRISPR array was present in the whole population, as shown by consistent read coverage. We infer that the lack of CRISPR targeting by the majority of the I57_Bv_host population members enabled the presence of I57_Bv_phage in W1 and W6.

**Figure 3.**
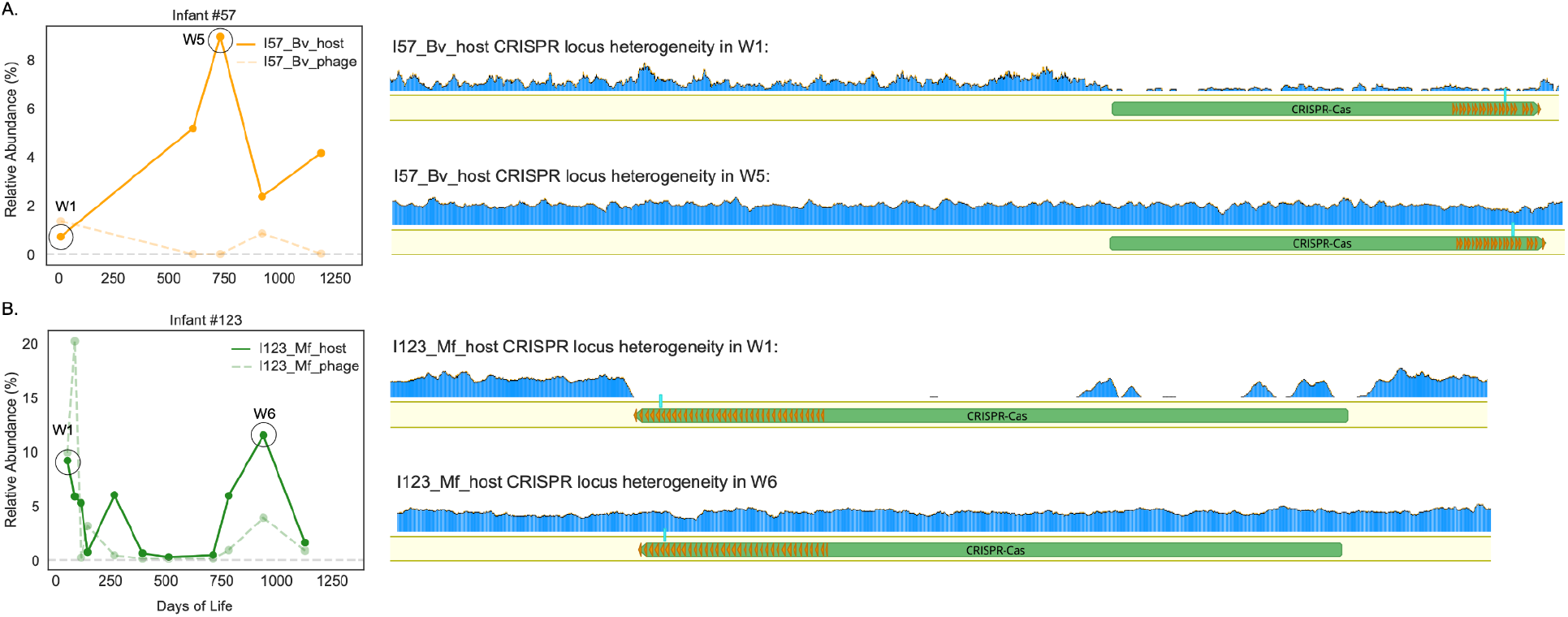
Co-persistence of phage and its predicted bacterial host. Relative abundances of I57_Bv_phage-I57_Bv_host (A) and I123_Mf_phage-I123_Mf_host (B) in infants #57 and #123, respectively, are shown as line plots on the left panel. For each infant, two bacterial CRISPR locus coverage files at two different time points are shown on the right panel as examples. CRISPR repeats are shown in orange arrows, and the spacer targeting the phage (I57_Bv_phage in panel A and I123_Mf_phage in panel B) is highlighted in aqua.

Unlike I57_Bv_phage, I123_Mf_phage was detected consistently throughout all sampling windows. Its host, *M. funiformis* (“I123_Mf_host”), initially did not encode the CRISPR array (during W1). However, starting in W2, the entire I123_Mf_host population encoded the CRISPR array targeting I123_Mf_phage (Figure 3B). One spacer targets a 36-nt region in phage’s tape measure protein. Prior to W5, the I123_Mf_phage region targeted by the CRISPR spacer did not accumulate any single-nucleotide polymorphisms (Figure S5A). Starting in W5, one SNP (C → T; leucine to phenylalanine nonsynonymous mutation) in the spacer-targeted region was detected in a sub-population, but this sub-population never completely overtook the original I123_Mf_phage population (Figure S5B-D). Interestingly, mapping of reads from some time points shows the same or lower coverage of the prophage compared to the flanking host genome. Reads from other time points show that the phage region coverage is higher than that of the flanking host genome (Figure S6). This indicates that this phage is periodically excised from the host genome and sometimes coexists in a non-integrated form (possibly as phage particles).

### Persisting phages in infants accumulate many SNPs in nucleic acid metabolism genes

Besides one phage persister (I57_Bv_phage) that was only found at the first and the last windows, the other 154 phage persister strains were detected in more than half of the time windows (Figure S3), indicating that the vast majority of persisters stably colonized the infant gut for three years. We hypothesized that some genes may have accumulated mutations to evade host defense systems and enable persistence. Thus, we quantified population-level single-nucleotide polymorphisms (pSNPs) in all genes encoded by persisting phages (Methods).

Out of 7365 genes encoded by phage persisters, ∼97% did not accumulate any fixed population-level mutations, whereas ∼3% (n=237) had at least one fixed pSNP. Of these, the majority had only one fixed pSNP, but 22 accumulated a significantly high length-normalized number of mutations over three years, with most mutations being synonymous (>1.5x interquartile range (IQR) of all non-zero mutation counts) (Table S5) (Methods). Of the 22 genes, 12 had no known function. Of those with annotations, genes involved in nucleic acid metabolism, regulation, and recombination were significantly enriched (p = 0.0216; Fisher’s exact test). The top five mostly mutated genes are a peptidase/endolysin, very late expression factor 1/tyrosine recombinase, reverse transcriptase (RT), a virion structural protein, and a protein of an unknown function (Table S5).

We further assessed mutations accumulated in maternal phage persister strains. Of the 28,538 genes examined, ∼2.8% had at least one fixed pSNP. Of these, 77 genes accumulated a significantly high length-normalized number of population-level mutations (Table S6), with most (n=52) having unknown functions. Interestingly, unlike infants, half of the mutations were non-synonymous. Of highly mutated genes that had annotations, no functional category was significantly enriched. The top five mostly mutated genes include three with no known functions, a peptidase/endolysin, and a thioredoxin (“thioredoxin_4”) (Table S6).

### Phages with reassigned stop codons persisted in infants and mothers

While ∼99.7% of the gut phageome of infants and mothers used standard genetic code (code 11), a small fraction of phages recoded the TAG or TGA stop codon to encode glutamine or tryptophan (genetic codes 15 and 4, respectively). We identified 37 recoded phage strains from 24 infants and 41 recoded phages from 22 mothers, most of which recoded the TAG stop codon (n=69). While the majority of the recoded phages were predicted to be lytic, five were temperate. Genome sizes of recoded temperate phages (28.6 ± 3.7kb; 100% CheckV complete) were significantly smaller than those of lytic phages (124 ± 35.6kb) (p = 0.0011; Wilcoxon rank-sum test). Recoded phages, on average, colonized infants for 146 days (∼5 months), and the majority of them appeared in infants after age one (Figure S7). Approximately half of the recoded phages found in infants were maternally transmitted.

Recoded phages were significantly more likely to persist in infants than phages using standard genetic code (p = 0.010; Fisher’s exact test). Ten recoded phages colonized infants during W1, and three (I57_PsAC1, I79_PsAC1, and I123_PsAC1), all using genetic code 15 and having a genome length of 99.6 ± 1.5kb, persisted until age three. These were from biologically unrelated infants #57 (preterm), #79 (full-term), and #123 (full-term), respectively. *Bacteroides vulgatus* was predicted to be the host for I57_PsAC1 using CRISPR spacer matches, but the hosts for the other two phages are unknown (Methods). All recoded persisters were maternally transmitted and persisted in mothers as well (Methods). Interestingly, these recoded persisters were all predicted to be lytic. Their stable gut colonization thus motivated us to seek genomic traits that may enable their persistence.

For all three phages, genes that contained in-frame TAG were significantly more likely to be mutated than genes lacking in-frame TAG (p = 5.01e-08; Fisher’s exact test). Recoded genes also accumulated a higher number of SNPs per nucleotide than genes that were not recoded (p = 0.0036; Wilcoxon rank-sum test). This may simply reflect that fast evolution is the predictor of the appearance of reassigned codons within genes.

All three phages displayed different patterns of change in their in-frame TAG content over time. First, the infant-associated phage I57_PsAC1 had a 7.0% net increase in in-frame TAG, mostly as synonymous mutations in structural genes (Figure 4A). In some cases, in-frame TAG was accumulated as a result of indels (Table S7). For the same I57_PsAC1 phage in infant 57’s mother, we detected only a 2.2% increase in in-frame TAG, but this rate is higher than observed for the other two phages (Figure 4A). The second phage, I79_PsAC1, rarely had a change in in-frame TAG (Figure 4A). Specifically, after three years, the in-frame TAG content had decreased by 0.52%. In the third case, I123_PsAC1, we detected a ∼2% net decrease in in-frame TAG by year 3. This drop in in-frame TAG was mostly a result of a 502-nt deletion within the recoded reverse transcriptase (RT), a key component of diversity-generating retroelements (DGRs^47, 48^) (Figures 4C and S7C).

**Figure 4.**
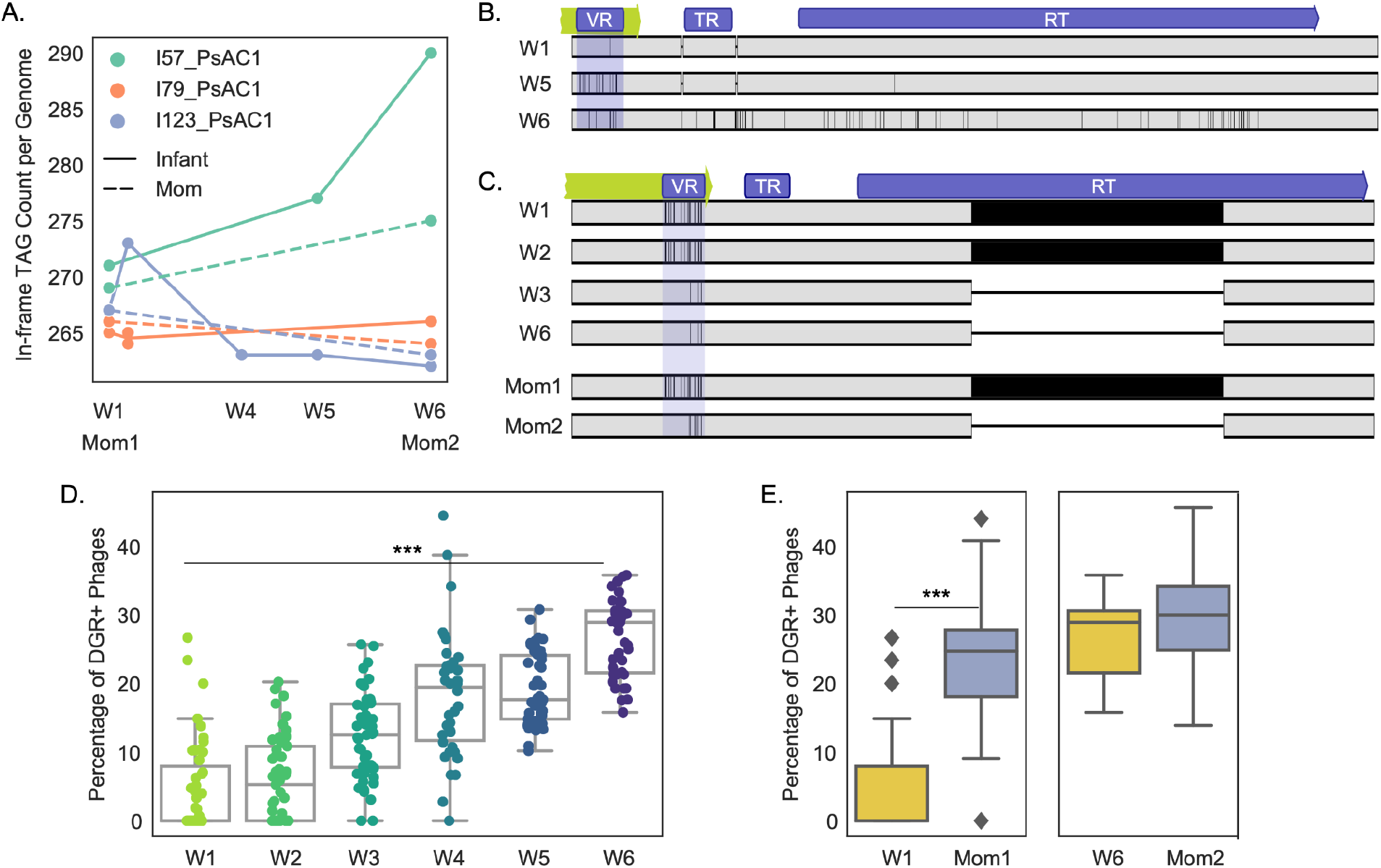
DGRs are associated with phage persistence. (A) The total number of in-frame TAG codons per persisting phage genome over time. Colors represent different persisting phage genomes. Infant persisters are shown in a solid line, and maternal persisters are shown in a dashed line. (B-C) Nucleotide alignments of DGRs encoded by phages I57_PsAC1 (B) and I123_PsAC1 (C) recovered over the three-year sampling period. Each row represents a time window-specific phage DGR region, and the solid vertical black lines represent SNPs that differ between sampling windows. (D) Percentages of DGR-encoding phages over time. Each dot represents an infant and is colored by infant age at the time of sampling (*** = p < 0.001). (E) Comparison of percentages of DGR-encoding phages in infants (gold) and mothers (cornflower-blue) during the initial (left panel) and the final (right panel) sampling windows (*** = p < 0.001).

### Diversity-generating retroelements may enable phage persistence in the gut

A DGR system was found in all three recoded persisters (I57_PsAC1, I79_PsAC1, and I123_PsAC1). As expected, all consist of an RT, a variable region (VR), and a template region (TR) (Figures 4 and S7; Table S8) (Methods). For each recoded persister, the predicted RT had multiple in-frame TAG codons and was adjacent to its predicted targeted gene, a recoded tail protein (Figures 4B-C and S7). Genomic alignments revealed that for each persister, the VR, which is located near the 3’-end of the targeted tail protein, mutated within the 3-year sampling period (Figures 4B-C and S7).

DGRs were present in I57_PsAC1 from the initial to the final sampling windows (Figures 4B and S7A; Table S8). For I79_PsAC1, despite the presence of an RT across all time points, a DGR was only predicted to be present during W6 and Mom2 (Figure S7B and Table S8). For I123_PsAC1, a 502-nt deletion within the RT during W3 rendered the DGR non-functional, and the VR of the targeted tail protein on I123_PsAC1 remained largely the same over the remaining time windows (Figures 4C and S7C; Table S8) (Methods). The same phage in the mother acquired the same indel in the RT (Figures 4C and S7C). Given that the VR of I123_PsAC1 was identical in the infant and the mother prior to W2 but differed by year 3 (W6 and Mom2), we speculate that the loss of DGRs might have occurred independently in the infant and the mother.

DGRs found in phages often target host-recognition regions such as tail proteins, enabling host tropism^47–50^. The identification of DGRs in all three recoded persisting phages, as well as the accumulation of a high number of mutations in persisting phages, regardless of genetic code, led us to hypothesize that gut phages may use DGRs for long-term persistence. Indeed, we found that persisting phages in both infants and mothers were significantly more likely to encode DGRs than non-persisting phages (p = 4.94e-27; Fisher’s exact test). Further, we observed that the percentage of DGR-encoding phages in infants increased over time (Spearman correlation r = 0.76, pvalue = 4.06e-53) (Figure 4D). Moms initially had a higher percentage of DGR-encoding phages than infants (q = 1.56e-12; Wilcoxon rank-sum test) (Figure 4E). However, by age three, the density of DGR-encoding phages was comparable between infants and mothers (q = 0.092; Wilcoxon rank-sum test) (Figure 4E).

## Discussion

We conducted strain-resolved analyses using a *de novo* constructed phage database to investigate phageome succession in preterm and full-term infants during their first three years of life. The inclusion of maternal fecal samples that were collected three years apart enabled us to assess the influence of the maternal phageome on early-life phageome assembly. The longitudinal maternal sample collection also enabled us to explore viral persistence in mothers. Our study differs from prior infant virome studies^29, 30, 33^ by resolving phageome succession with strain resolution, enabling us to differentiate initially colonizing phage strains from phages that were acquired later in time. Enabled by the three-year sampling period for both infants and mothers, we explored the topic of within-individual phage succession by primarily focusing on phage early colonizers that persisted for nearly three years in infants and mothers.

The infant gut phageome underwent a major phage population turnover between birth and age three, yet approximately 9% of initial colonizing phages persisted over this time period. The maternal gut phageome was identified to be a critical source of these long-term persisting phages. Phage strains that were vertically transmitted were more likely to persist in both mothers and infants than those that were not transmitted, implying continuous and reciprocal seeding between mothers and infants. Colonization by maternally transmitted phages could also be a result of the co-transmission of their bacterial hosts, such as *Bacteroides*, a commensal typically colonizes the gut long-term^13, 51^. Further, the persistence of these maternally transmitted phages could reflect their adaptation to the gut environment, such as better adherence to the gut mucosal^21, 52^ or tolerance by and/or to the immune system^53–56^.

*Bacteroides* was the most likely genus to harbor persisting phages. Bacterial strains of this genus were also more likely to persist in infants than strains of other genera. Bacterial persistence requires mechanisms, such as using CRISPR-Cas systems^57, 58^, to evade phage predation, which often results in the killing of phages. The observation of bacteria-phage co-persistence suggests a compromise might have been reached between the two entities. We observed that while some *B. vulgatus* have a CRISPR-Cas system that targets the persisting *B. vulgatus* phage, the majority of the population lacks this locus, resulting in a mix of phage-sensitive and phage-resistant bacterial host strains. We reasoned that the presence of mixed host populations with varying degrees of phage sensitivity may have ensured prolonged phage-bacteria co-existence. Indeed, similar phenomena had been seen in *Bacteroides* spp. with phase-variable polysaccharides, leaving only a subset of host populations that are sensitive to phage infection^24, 59, 60^.

Another factor that may contribute to phage persistence is the existence of a diverse pool of phage populations. We found that phage persisters had a significantly higher nucleotide diversity than non-persisters, implying that they may be more likely to evade bacterial defense systems when compared to non-persisters. Supporting this hypothesis is the observation that diversity-generating retroelements (DGRs) were more likely to be encoded by persisters than non-persisters during the initial colonization period. DGRs are often used by phages to diversify certain genomic regions, many of which are host-recognition regions^47–50, 61^. DGRs were present in all three recorded phage persisters, and time-dependent tail protein diversification was seen in all of them. We speculate that active tail protein diversification, which enables adaptation to their host’s evolved receptor, is one strategy used by these phages to prolong their stay in the gut. In addition, tail protein modification may enable binding to a new host receptor, altering the phage host range. In the human gut, where microbial cells are densely populated and strain heterogeneity is high^62–65^, being able to infect multiple hosts is suggested to be advantageous for long-term phage colonization^34, 66^.

Despite constituting a small percentage of the gut phageome, early colonizing phages that use the reassigned TAG stop codon were significantly more likely to persist than phages that used a standard genetic code. The re-coding of TAG to incorporate glutamine is a phenomenon that appears to emerge periodically, given lineages where closely related phages adopt the standard and/or TGA-reassigned genetic codes^67^. Time-series population genetic analyses revealed evidence of in-frame TAG accumulation in both infants and mothers over time. However, for the three re-coded persisting phages, there was a large variation in the number of in-frame TAG codons over three years, which may imply different lengths of time since they adopted alternative coding. It is possible that differences in the number of in-frame TAGs simply reflect selection for different populations rather than *in situ* evolution, and this may explain the case where the in-frame TAG content both increased and decreased (phage I123_PsAC1). However, in one case (phage I57_PsAC1), the consistent increase in in-frame TAGs at every sequential time point is very likely to be the result of ongoing in-frame TAG introduction. For this re-coded *Bacteroides* lytic phage, we observed a significant expansion of in-frame TAG in primarily “late” structural and lysis genes in a preterm infant and their mother over the three-year sampling period. Introducing in-frame TAG in late-expressed genes was recently proposed as a way for re-coded phages to prevent premature lysis^67, 68^. The expansion of in-frame TAG may be a result of phages counteracting bacterial immune systems that force early lysis^69–72^. Given the initially low bacterial diversity of the infant gut microbiome^6, 8, 10, 13^, we speculated that genomic recoding might be particularly beneficial for certain lytic phages to establish long-term colonization since it likely provides the phage with another layer of defense against bacterial immune systems. Interestingly, the persisting recoded *Bacteroides* phage in an infant accumulated many more in-frame TAGs than did the phage in the mother. It is hard to offer an explanation for this, but the observation suggests a greater need for a competitive advantage for phages in the rapidly changing infant gut microbiome.

We also found phage persistence to be affected by prematurity. Compared to full-term infants, preterm infants tend to have initial gut microbiomes that are populated by hospital-associated strains, and undergo more drastic microbiome shifts in order to reach a stable, full-term-like state^13^. We previously reported that full-term infants were more likely to be colonized with persisting bacterial strains than preterm infants^13^. In the current study, we noted that full-term infants have more persisting phages than preterm infants. The lack of long-term persisting phage in preterm infants could be partly attributed to the lack of early-colonizing *Bacteroides* strains^13^, the genus that was found to be enriched with harboring persisting phages in our study here.

We observed an increased viral diversity as infants mature. Similar to the trend seen in gut bacterial communities, we found the gut phageomes of both pre-and full-term infants reached a comparable level of complexity as those of mothers after three years. Our findings are in contrast with two previous early-life virome studies that reported a decrease in viral diversity over two to three years. The study by Lim and colleagues characterized their viromes primarily using the NCBI nucleotide database that was updated to 2013^30^. The study led by Walters and colleagues relied on the NCBI nucleotide database that was updated to 2019^33^. These databases contain far fewer gut phage sequences than databases generated over the years following^42, 43, 45, 73, 74^, thus may have missed a significant number of phages present in individuals and not represented by public database sequences. In addition, these studies performed sequencing on enriched virus-like particles (VLPs), rather than on the whole metagenomes like we did here, thus may have missed nearly all prophages. Given the high individuality and turnover rate of infant gut phageomes^73, 75^ and incomplete public gut phage databases^31, 46, 76–78^, we reason that use of a *de novo* constructed, study-specific phage database may have provided a more precise and representative picture of the early-life gut viral assembly.

In summary, by assessing persisting phages in infants and their mothers, and evaluating factors associated with long-term phage colonization, our study provides a fine-grained view of the early-life gut phageome succession. Despite consisting of a small fraction of the initial phage colonizers, phage persisters likely play a significant role in shaping the developmental trajectory of the infant gut microbiome. By identifying and tracking individual phage and bacterial strains in preterm and full-term infants, as well as their mothers, through the first three years of life and through population genetics analyses, we determined that maternal origin, the persistence of bacterial hosts, a high population diversity, and genetic recoding all contribute to phage persistence in the human gut.

## Lead Contact

Further information and requests for resources should be directed to the Lead Contact, Jillian F. Banfield.

## Acknowledgments

We thank Rohan Sachdeva, Jordan Hoff, and Shufei Lei for their technical support. We are also grateful for all the families that participated in this study. For funding support, we acknowledge NIH award RAI092531A to J.F.B and M.J.M.

## Author Contributions

Y.C.L., M.J.M., and J.F.B. designed the study; B.A.F. performed DNA extractions of fecal samples; Y.C.L. coordinated the acquisition of and performed analysis on the metagenomics data; Y.C.L. and A.B. constructed the phage genome database; LX.C. assisted in phage data analyses; Y.C.L and J.F.B. wrote the manuscript and all authors contributed to the manuscript revisions.

## Data and Code Availability

Metagenomics sequencing reads, metagenome-assembled bacterial and phage genomes will be deposited on NCBI soon.

## Methods

### Study Details

This study was reviewed and approved by the University of Pittsburgh Human Research Protection Office (IRB STUDY19120040). This nested case-control observational study was originally designed to study the gut microbiomes of premature and full-term infants as well as the gut microbiomes of premature infants who developed NEC and/or LOS and age-matched premature infants over the first year of life. For these purposes, we enrolled a total of 183 infants (35 full-term infants and 148 preterm infants born before 34 weeks of gestation). Please refer to Lou et al.^13^ for more details on infant enrollment. Ultimately, we acquired longitudinal samples from 28 full-term and 24 preterm (9 healthy controls, 9 NEC infants, 5 LOS infants, and 1 infant that developed both NEC and LOS) from birth to up to age three (Figure S1).

Fecal samples from enrolled infants and their mothers were all collected at the UPMC Magee-Womens Hospital (Pittsburgh, PA) over the course of five years. While full-term infants were discharged from the hospital within 3 days after birth and received no perinatal antibiotics, all preterm infants received empiric antibiotics immediately following birth during an evaluation for early-onset sepsis and then spent their first 2 to 3 months in the hospital. In addition to infant fecal samples, we collected up to two fecal samples from 46 mothers of 52 infants, one within the first two weeks after delivery and one when infants turned three years old (Figure S1). All samples were collected with parental consent, and subjects were de-identified before the receipt of samples. De-identified metadata for all 52 infants and their mothers were provided in Table S1.

### Sample collection and metagenomic sequencing

Infant and maternal fecal samples were collected either at UPMC Magee-Womens Hospital by trained nurses or at home by parents provided with detailed collection instructions. Specifically, fresh infant stool samples were collected directly from infants while they were actively excreting or from diapers shortly after the stools were released. Maternal fecal samples were collected using a commode specimen collector, from which fecal samples were transferred into a collection tube. All stool samples collected at the hospital were immediately stored at −80°C following collection. Samples collected at home were stored in home freezers until they were picked up by research staff and transferred to the −80°C condition. DNA extraction of frozen fecal samples collected when infants were two years old or younger was performed via the Qiagen DNeasy PowerSoil HTP 96 DNA isolation kit with modifications to the manufacturer’s protocol (plate-based extractions; a total of 9 plates were used for 702 fecal samples). DNA extraction of fecal samples collected post age-two was performed using the Qiagen QIAamp PowerFecal Pro DNA Isolation Kit (single-tube extractions; used for 117 samples). For each 96-well extraction plate, at least one reagent-only negative control was included. Two ZymoBIOMICS Microbial Community Standards (catalog no. D6320 and D6321) were also included as positive controls for four DNA-extraction plates.

Metagenomic sequencing of collected infant and maternal fecal samples was performed in collaboration with the California Institute for Quantitative Biosciences at UC Berkeley (QB3-Berkeley). Library preparation on all samples was performed as previously described ^79^. Final sequence-ready libraries were pooled into subpools and visualized and quantified on the Advanced Analytical Fragment Analyzer. All libraries were then evenly pooled into a single pool and checked for pooling accuracy by sequencing on Illumina MiSeq Nano sequencing runs. The single pool was adjusted based on the MiSeq sequencing run and sequenced on individual Illumina NovaSeq6000 150 paired-end sequencing lanes with 2% PhiX v3 spike-in controls. Post-sequencing bcl files were converted to demultiplexed fastq files per the original sample count with Illumina’s bcl2fastq v2.20 software.

### Metagenomic assembly and gene prediction

Reads from all 819 samples were trimmed using Sickle (https://github.com/najoshi/sickle), and reads that mapped to the human genome with Bowtie2^80^ under default settings were discarded. Reads from each sample were then assembled independently using IDBA-UD^81^ under default settings. Co-assemblies were also performed for each infant, in which reads from all samples of that infant were combined and assembled together. Scaffolds that are <1 kb in length were discarded. The remaining scaffolds were annotated using Prodigal^82^ to predict open reading frames (ORFs) using default metagenomic settings. tRNAs were predicted using tRNAscan-SE (v0.1)^83^.

### Microbial metagenomic *de novo* binning

Pairwise cross-mapping was performed between all samples from each individual to generate differential abundance signals for binning. Each sample was binned independently using three automatic binning programs: metabat2^84^, concoct^85^, and maxbin2^86^. DasTool^87^ was then used to select the best microbial bins from the combination of these three automatic binning programs. The resulting draft genome bins were dereplicated at 98% whole-genome average nucleotide identity (gANI) via dRep^88^ (v3.4.3) *dereplicate* (-comp 75 -con 10 -sa .98 -nc .25). Genomes with gANI ≥98% were classified as the same bacterial subspecies, and the genome with the highest dRep score was chosen as the representative genome from each bacterial subspecies, resulting in a total of 1,951 de-replicated microbial subspecies.

### Taxonomy assignment

The amino acid sequences of predicted genes of all assembled bins were searched against the UniProt100 database using the usearch ublast command with a maximum e-value of 0.0001. tRep (https://github.com/MrOlm/tRep/tree/master/bin) was used to convert identified taxIDs into taxonomic levels. Briefly, for each taxonomic level (species, genus, phylum, etc.), a taxonomic label was assigned to a bin if ≥50% of proteins had the best hits to the same taxonomic label. GTDB-Tk^89, 90^ (v2.1.1) was used to resolve taxonomic levels that could not be assigned by tRep.

### Phage prediction

Phage prediction tools Seeker^91^, VIBRANT^92^, and geNomad^93^ were run on assembled metagenomes (contigs ≥ 4.5kb) using default settings. Assembly-free-based phage prediction was performed using Phanta^46^ on all trimmed, human-DNA-removed reads under default settings. Prophages were identified and trimmed by removing flanking host regions using VIBRANT, geNomad, and CheckV^94^. Free-existing linear phage fragments were extended using COBRA^95^. CheckV was run on all predicted phages and trimmed proviruses to evaluate completeness and quality. Contigs evaluated as low quality by both CheckV and VIBRANT and had a geNomad viral score <0.9 with ≤1 geNomad viral hallmark gene were removed from the analysis. Contigs with eukaryotic viral taxonomies assigned by geNomad and/or CheckV were also removed. In total, 32,401 phage scaffolds were generated, and they were mostly medium-and high-confidence phages with (1) contigs < 100 kb with viral genes > host genes, (2) contigs > 100 kb with < 20% host genes, or (3) host-region-trimmed prophages with a minimal length of 25 kb.

To generate a non-redundant phage reference database, all 32,401 phage scaffolds were dereplicated at 98% gANI over 85% of the phage genomes using dRep *dereplicate* (-sa 0.98 --ignoreGenomeQuality -l 4000 -nc 0.85 --clusterAlg single -N50W 0 -sizeW 1). Genomes with gANI ≥98% were classified as the same phage subspecies, and the phage genome with the highest dRep score was chosen as the representative genome from each subspecies, resulting in a total of 9,929 phage subspecies. Manual inspections using phage gene annotations (see “Phage code prediction and gene annotations”) additionally removed contigs that were 1) evaluated as low quality by CheckV, VIBRANT, or geNomad (viral score < 0.9) and 2) lacked phage hallmark genes (i.e., phage structural genes), resulting a final set of 8,424 representative high-confidence phage subspecies.

VIBRANT was used to predict the lifestyle of all free-existing phages, and all trimmed prophages were assigned as temperate phages.

### Identification of novel phage species

Reference phage sequences retrieved from the five studies^31, 42–45^ were clustered with our reconstructed 8,424 phage genomes at 95% ANI over 85% of the length. Genome clustering was performed using the greedy, centroid-based algorithm developed by Nayfach et al. (see the “supporting code” section at https://bitbucket.org/berkeleylab/checkv/src/master/)^96^. We chose 95% ANI because numerous studies have suggested that this threshold groups closely related and biologically relevant phages into “viral species”^31, 42, 97, 98^.

### Phage code prediction and gene annotations

Identification of stop-codon reassigned phages was performed on all predicted phage genomes as previously described^67^. Coding sequences were subsequently generated using Prodigal, using genetic code 4 for TGA-recoded phages, code 15 for TAG-recoded phages, and code 11 for remaining standard-code phages. HMMER^99^ was used to annotate the resulting sequences with the PFAM, pVOG, VOG, and TIGRFAM HMM libraries. In some cases, proteins were annotated by running BLASTP searches against the NCBI database. To further annotate genes with no known functions, all phage proteins were clustered into protein families created using a two-step protein-clustering method^100^. Proteins were first clustered into subfamilies using MMseqs^101^, and HHBlits^102^ was used to generate HMMs of each subfamily based on alignments generated with the MMseqs result2msa parameter. The resulting HMMs were further compared to one another using HHBlits, and MCL clustering was used to generate families from the HMM-HMM comparisons. All annotations were merged and the annotation with the lowest e-value was chosen. Functional categories were assigned using the modified annotation sheets published by Pfeifer and colleagues^103^.

### Phage host prediction

Hosts for prophages were assigned based on the genome into which the contig containing prophage was binned. The resulting taxonomy was further verified via taxonomic profiling. Specifically, contig-based taxonomic profiling was performed by using DIAMOND^104^ (fast mode, e = 0.0001) to search all phage and bacterial proteins against a custom version of the UNIREF100 database that retained NCBI taxonomic identifiers. tRep was then used to profile the taxonomy of each contig. For each contig, the microbial taxonomy with the most hits was considered to be the putative host, but only if that taxonomy had more than 3x hits than the second most common taxonomy^67, 105^.

For prophage-containing contigs that were not assigned a genome bin, as well as free-existing phage scaffolds, a combination of CRISPR spacer analysis and taxonomic classification was used to predict putative host taxonomy. Metagenome-specific CRISPR spacers were mined using minCED^106^ and PILER-CR^107^. The resulting 1,950,165 spacers were merged with >11 million spacers from CRISPRopenDB^108^ to generate a relatively comprehensive CRISPR spacer database with 13,717,947 spacers. Subsequently, *blastn-short* was run on the constructed spacer database to identify matches between phage and spacer with >90% ANI and >90% spacer coverage. In almost all cases (especially up to the host genus level), the CRISPR spacer analysis and the taxonomic profiling agreed on the phage host taxonomy. In cases where these two analyses were not in agreement, if the phage was targeted by ≥3 CRISPR spacers from ≥3 bacterial genomes with near-perfect spacer matches (≤2 mismatches) and if all these bacterial genomes shared the same taxonomy, the host taxonomy was then assigned using the consensus result of the CRISPR spacer analysis. Otherwise, the host taxonomy was considered unknown.

### Detection of subspecies, relative abundance calculation, and identification of bacterial and phage strains

Reads from each individual fecal sample were mapped to all concatenated dereplicated, representative phage (n=8,424) or bacterial (n=1,951) subspecies genomes (generated via dRep *dereplicate* as described above) using Bowtie2 under default settings. inStrain^109^ (v1.6.3) *profile* was run on all resulting mapping files using a minimum mapQ score of 0 and insert size of 160. Phage and bacterial genomes with ≥0.75 and ≥0.5 breadth, respectively, in samples were considered to be present, and their relative abundances were calculated as the percentage of total sample reads mapping to each genome, which were. Calculated relative abundances of phages and bacteria were subsequently normalized by the sum of relative abundances of all phages and bacteria, respectively, from each sample.

To identify near-identical phage and bacterial strains, inStrain *compare* was run to compare the genome similarity among all genomes that were present in ≥2 samples. Specifically, inStrain *compare* was used under default settings to compare read mappings to the same genome in different pairs of samples. Samples were considered to share the same phage or bacterial strain of the examined genome if the compared region of the genome from samples shared ≥99% or ≥99.999% population-level ANI (popANI), respectively. Only genomic areas with at least 5x coverage in samples were compared, and sample pairs with less than 75% or 50% of comparable regions of the phage or bacterial genome, respectively, were excluded.

### Detection of mother-to-infant vertical transmission

For each mother-infant dyad, every fecal sample from the infant was compared to their mother’s fecal samples using inStrain *compare* (described above) to search for identical strains of phages (≥99% popANI & >0.75 percent_genome_compared) or bacteria (≥99.999% popANI & ≥0.5 percent_genome_compared). A strain was considered to be vertically transmitted if it was shared between at least one maternal fecal sample and at least one infant fecal sample.

### Three-year persister and non-persister detection

“Beginning-end” and “pairwise” approaches were used to differentiate persisters from non-persister strains among early colonizers. The “beginning-end” approach searched for strains that shared ≥99% (phage) or ≥99.999% (bacteria) popANI between the first (≤month 2; W1) and the last sampling windows (≥month 30; W6). The “pairwise approach” identified strains that shared ≥99% (phage) or ≥99.999% (bacteria) popANI across ≥50% of the consecutive sampling windows (W1-W2, W2-W3, W3-W4, W4-W5, and W5-W6).

### Quantifying single-nucleotide polymorphisms (SNPs) accumulated in genes encoded by phage persisters over time

For all phage persisters, inStrain *profile* was used on default settings to identify single-nucleotide polymorphisms (SNPs) between all sample pairs. Only SNPs found within ORFs were retained for further analysis. To estimate fixed mutations over three years, phage genomes from the initial (W1 or Mom1) and the final (W6 or Mom2) sampling time points were compared. Genes accumulating a higher number of mutations than expected were identified using the interquartile range (IQR) outlier approach. Specifically, after normalizing for gene lengths, highly mutated genes were identified as those that accumulated SNP counts that were more than 1.5x IQR of all genes with ≥1 SNP.

### DGR-detection and time-series comparative genomic analysis

*DGR_identification* scripts (v1.0) (https://bitbucket.org/srouxjgi/dgr_scripts/src/master/)^47^ were run on all 32,401 predicted phage genomes to identify diversity-generating retroelements (DGRs). Target genes of DGRs were annotated using the PFAM, pVOG, VOG, and TIGRFAM HMM libraries as well as via BLASTP searches against the NCBI database (described above).

To examine DGR-induced mutations over time for a phage persister, fecal metagenome-specific phage genomes from the same dRep cluster as the representative phage persister genome were extracted (Cdb.csv; generated when running dRep *dereplicate*, as described above). Subsequently, DGR regions from all sample-specific phage genomes from the same individual were aligned and visualized using Geneious (https://www.geneious.com). In the case of I123_PsAC1, a 502-nt deletion within the recoded RT was observed in genomes recovered starting W3, which corresponded to the absence of DGRs predicted by *DGR_identification* scripts. We further confirmed the absence of the RT since W3 by comparing RT’s conserved domains with (“MKRYNNLFDKVVSLDNLYLADKKARRNKSHRKDIIEFDKNKDELLLQLQKQLIEGKYVTSEYHTFIIKEPKERIIFKLPYYPDRIVHHAIMNILEPIWCSVFITNTYSCIKKRGIHKALYDVQSALKDKQNTVYCLKLDVRKFYPSIDHEILKQIVRKKIKDNKLLALLDGIIDSVEGVPIGNYLSQFFANLYLSYFDHWLKEDKAVKYYFRYADDMVILHSDKEYLRQLLDEIREQLGTLKLEIKSNYQIFRVEDRSISFVGYRIYHDYTLIRKNIKHKMCKKVAAMNKLKHMTYSEYRQQVCSHIGWMKHCNGINLLKKIIKYHQLIEYARSS*”) and without (“MKRYNNLFDKVVSLDNLYLADKKARRNKSHRKDIIEFDK NKDELLLQLQKQLIEGKYVTSEYHTFIIKEPKERINQKLKVIIRYSEQKIEVYPLQDIESITIIL*”) the deletion using NCBI’s Conserved Domain Database (CDD)^110^.

### Community diversity analysis

Since the earliest fecal sample was collected several days after birth for preterm infants and around the first month of life for full-term infants, all community-diversity analyses between the two infant groups were conducted in the same chronological-age time frame (thus excluding any preterm samples taken before month 1).

To measure the alpha diversity change of the gut phageomes, if not otherwise specified, a Wilcoxon rank-sum test was conducted to compare gut phageomes at different sampling windows. A module from scikit-bio (http://scikit-bio.org/) was used to calculate the Shannon Diversity Index (“skbio.diversity.alpha.shannon”). Richness was calculated by quantifying the number of detected phage subspecies in each sample.

### Two-group univariate comparisons

Statistical significance was calculated using Fisher’s exact test (as implemented using the Scipy module “scipy.stats.fisher_exact”), Wilcoxon rank-sum test (as implemented using the Scipy module “scipy.stats.ranksums”), and the binomial test (as implemented using the Scipy module “scipy.stats.binom_test”). All multiple comparisons were false discovery rate (FDR) corrected with a threshold of q < 0.05. Sample correlation was calculated using the Spearman rank-order correlation coefficient (as implemented using the Scipy module scipy.stats.spearmanr).

## Supplemental Figures

**Figure S1.**
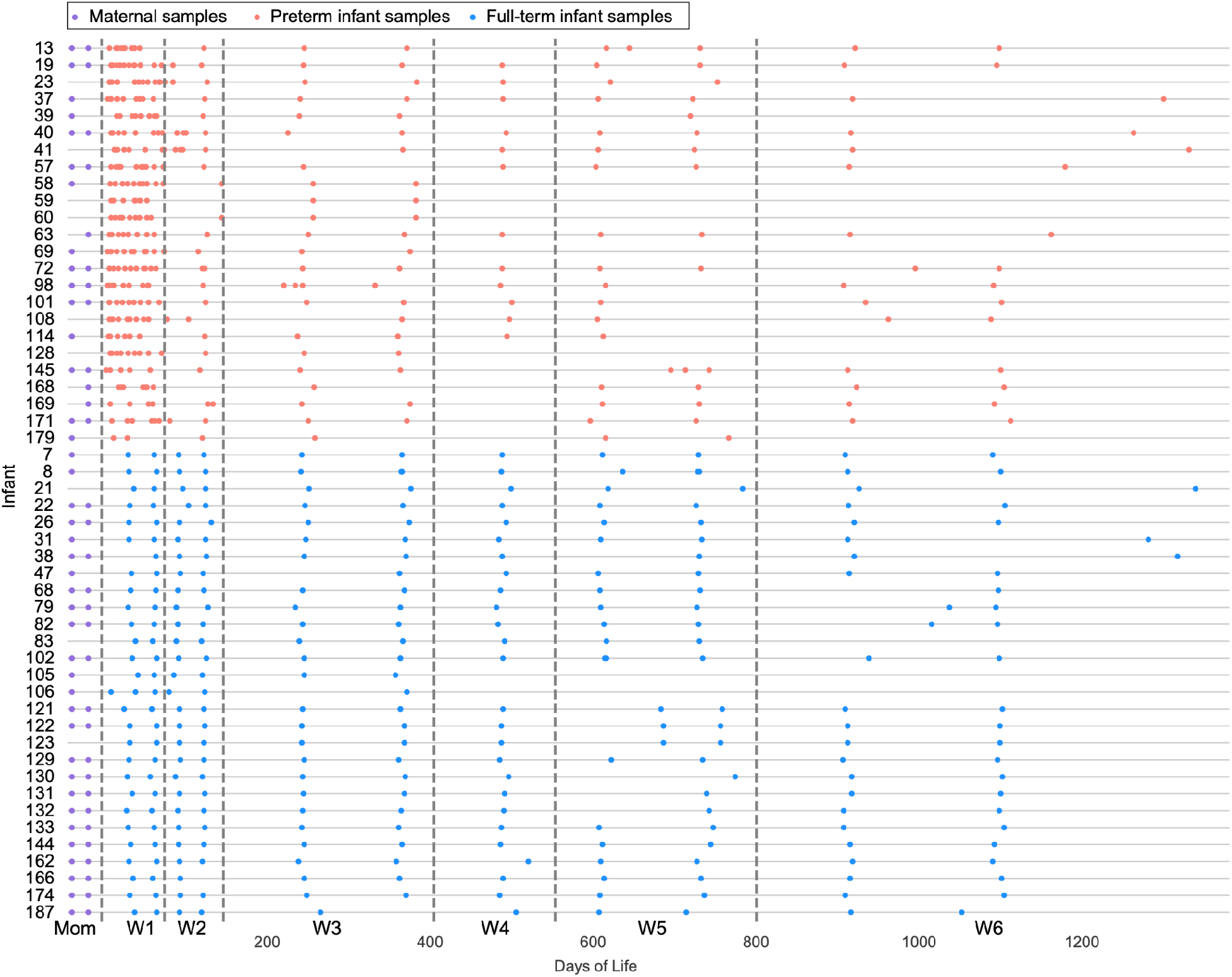
Fecal sample collection of all 52 infants. Each row represents an infant. Each solid circle corresponds to a sequenced fecal sample, and they are colored by the infant’s prematurity (salmon red: preterm infants; sky blue: full-term infants). Maternal fecal samples were collected around the time of delivery and when infants were three years old and are represented by solid purple circles before the first day of life of the matching infant. W1 to W6 represent the six time windows into which infant fecal samples were grouped.

**Figure S2.**
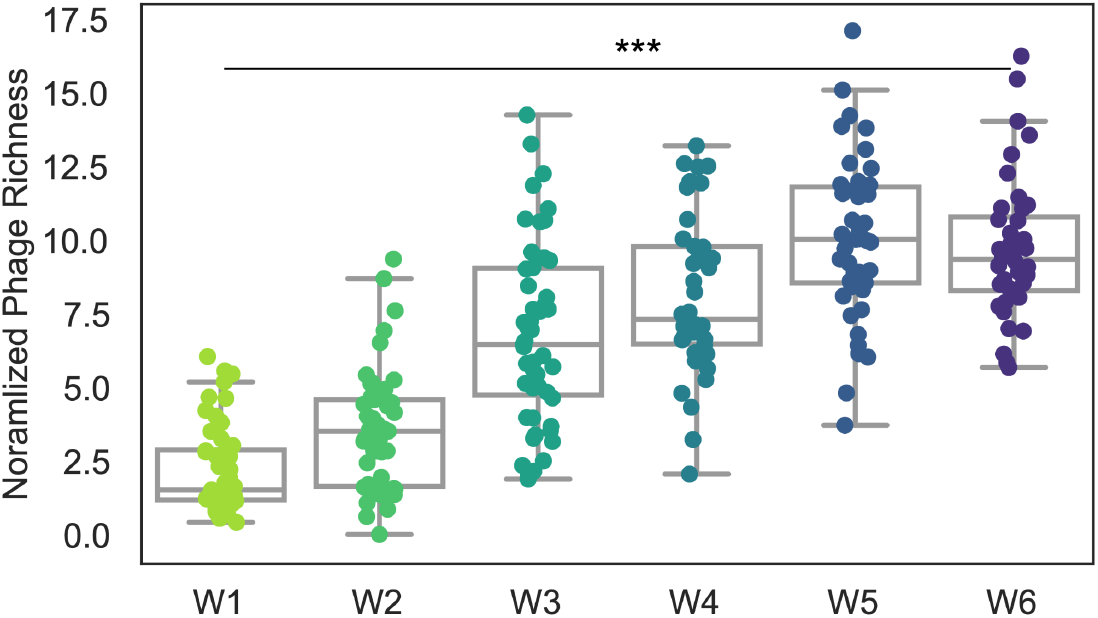
Alpha diversity of Phanta-characterized infant gut phages. The phage alpha diversity, measured via richness (normalized by sequencing depth), was quantified over time. Each dot represents an infant and is colored by infant age at the time of sample collection (*** = p < 0.001).

**Figure S3.**
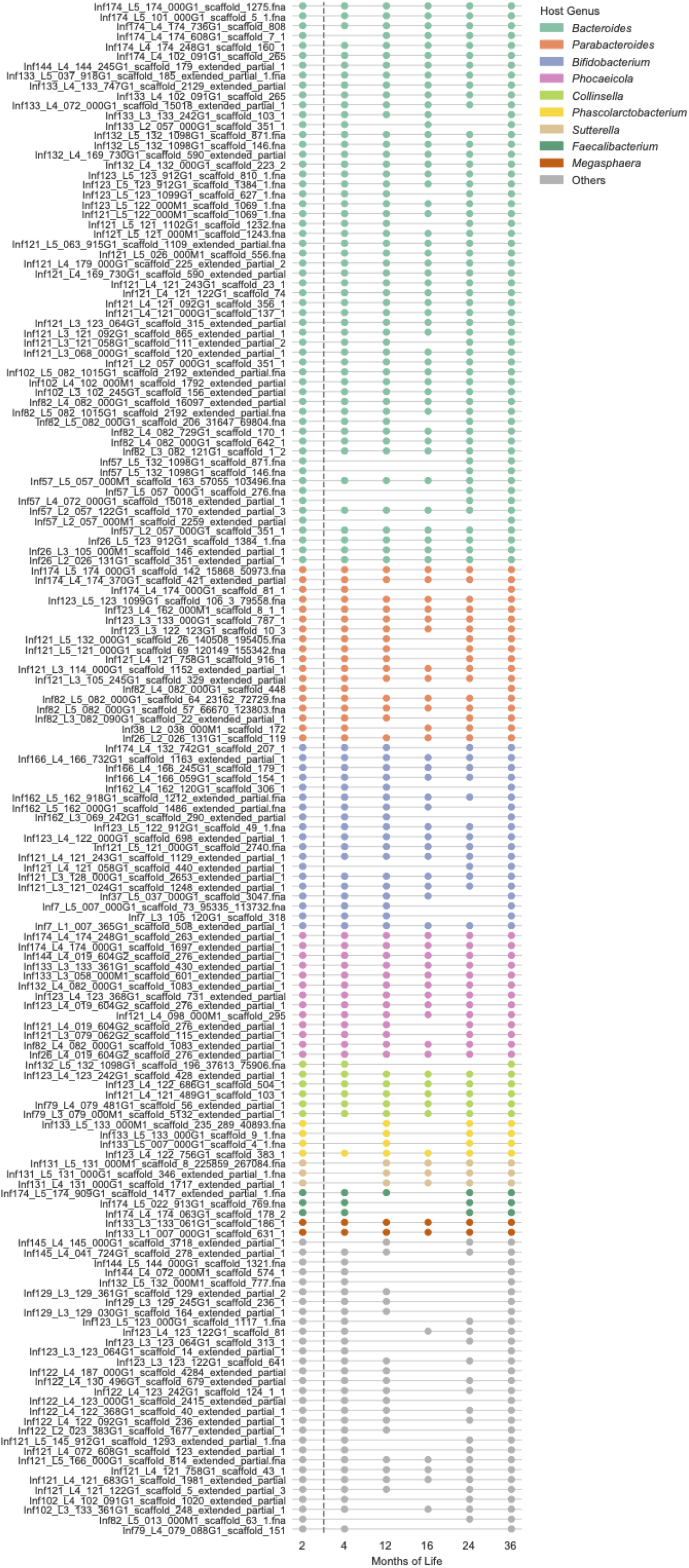
Infant gut phage persisters colonization overview. Schematic of all 155 infant gut persisting phage strains. Each row represents an infant-specific phage persister, and circles represent the months in which the strain was detected. Strains are colored by their putative bacterial hosts.

**Figure S4.**
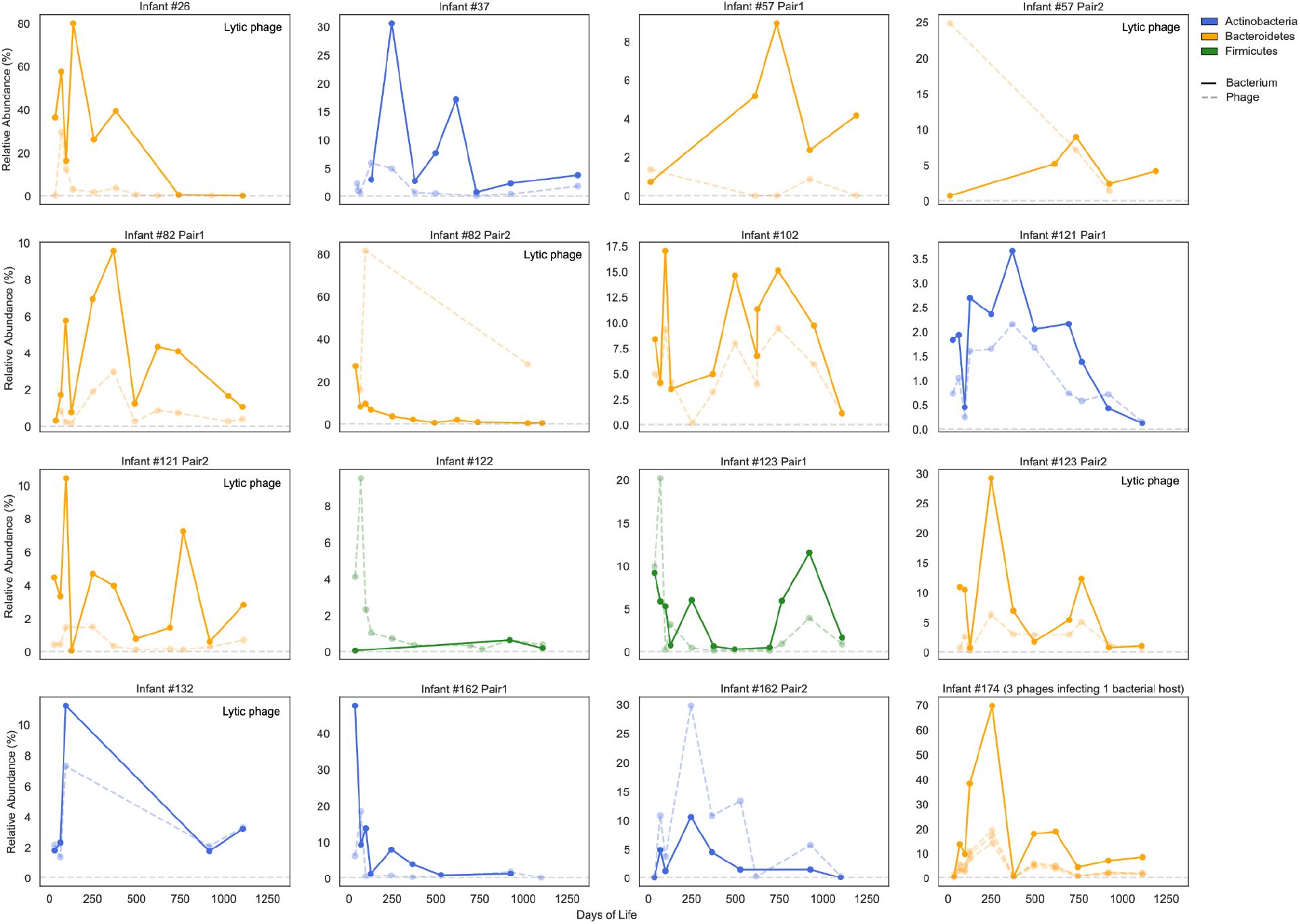
Co-persistence of phage and its predicted bacterial host strains. The normalized relative abundance of phage persister (in a dashed line) and its predicted bacterial host (in a solid line) over time. Lytic phages are noted in the subplots. Otherwise, phages are temperate. Lines are colored by bacterial host phyla. Except for the last subplot (at the lower right corner), the rest of the subplots all show the co-persistence of one bacterium-phage pair. In the last subplot, three phages were predicted to infect one Bacteroidetes genome; thus, they were all included in one subplot. Given the low abundance of phage in comparison to its bacterial host counterpart, we normalized the relative abundance of phages and bacteria separately. For instance, if a phage’s normalized relative abundance is 20%, it indicates that the phage represents 20% of the predicted phage population in the given sample.

**Figure S5.**
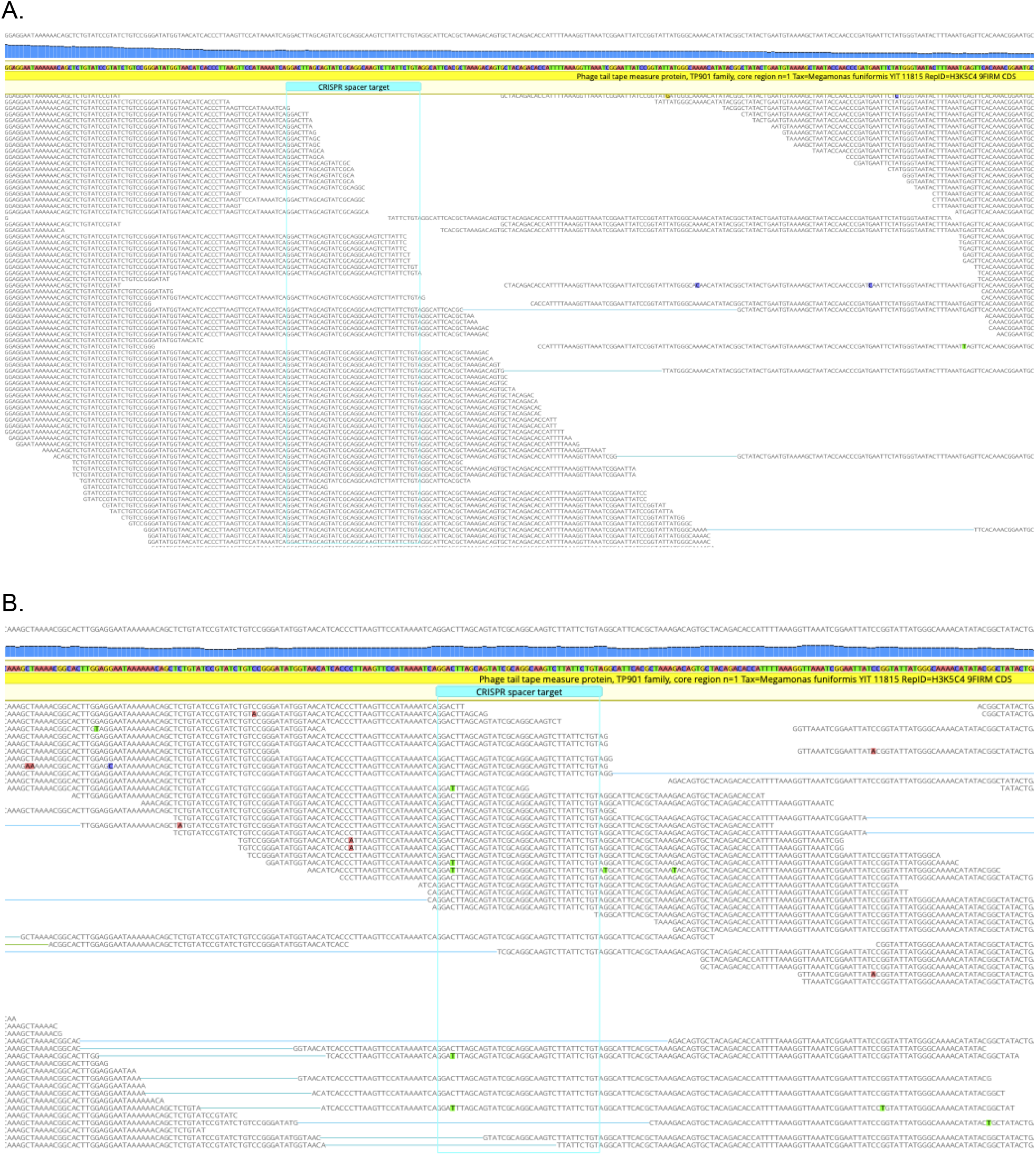
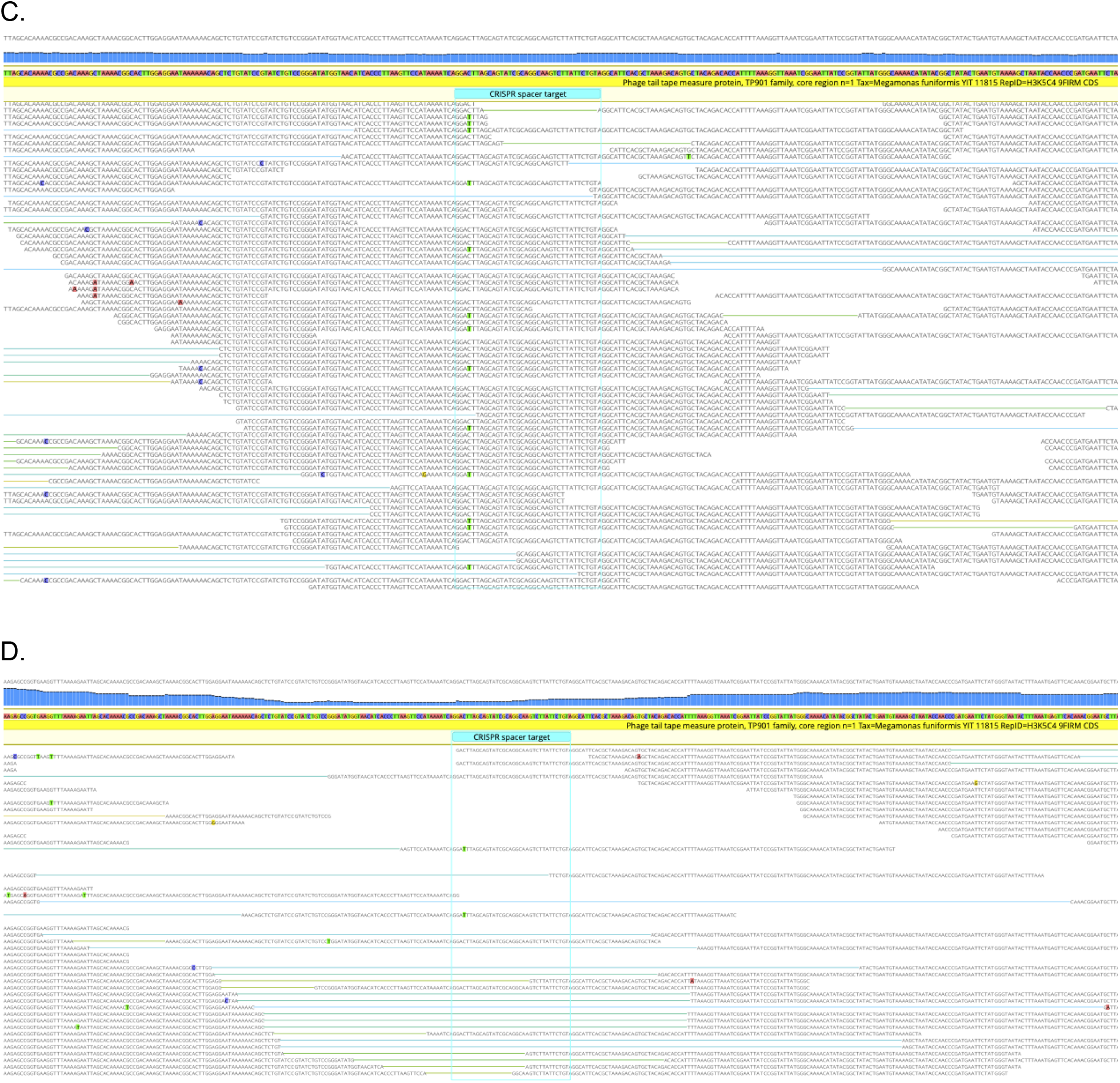
CRISPR spacer targeted region in I123_Mf_phage over time. (A-D) Reads from W1 (A), W5 (B), W6 day-of-life 912 (C), and W6 day-of-life 1099 (D) were mapped to the CRISPR targeted region of the I123_Mf_phage genome. An SNP (colored green; C → T) was detected in a subpopulation of the phage starting on W5. Figure was generated via Geneious.

**Figure S6.**
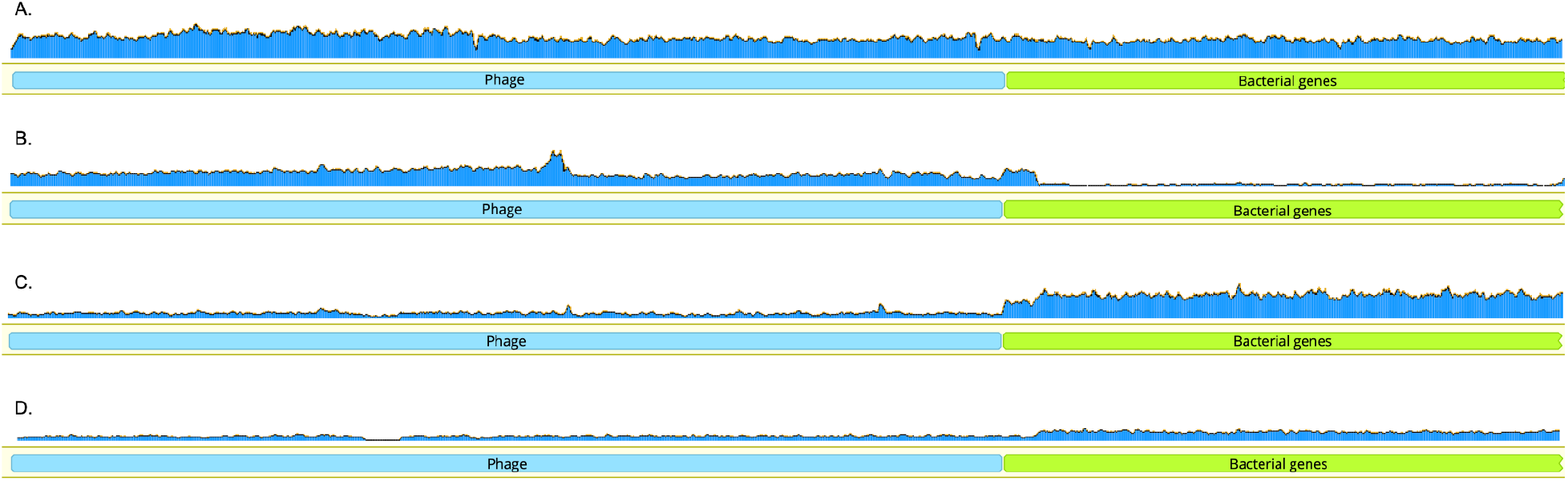
Coverage of the I123_Mf_phage over time. (A-D) Reads from W1 (A), W2 (B), W3 (C), and W5 (D) were mapped to the lysogenic contig in which the I123_Mf_phage was integrated into the host chromosome. I123_Mf_phage was considered actively replicating if the coverage of the phage is higher than that of bacteria, as seen in panel B, suggesting that the majority of the phage existed as free-existing particles. If phage’s coverage is the same as its host, as seen in panel A, it suggests that the phage exists as a prophage and nearly all bacterial population is lysogenized. For panels C and D in which the coverage of phage is less than that of bacteria, it suggests the partial lysogeny of the bacterial population and the phage likely exists as a prophage form only. Figure was generated via Geneious.

**Figure S7.**
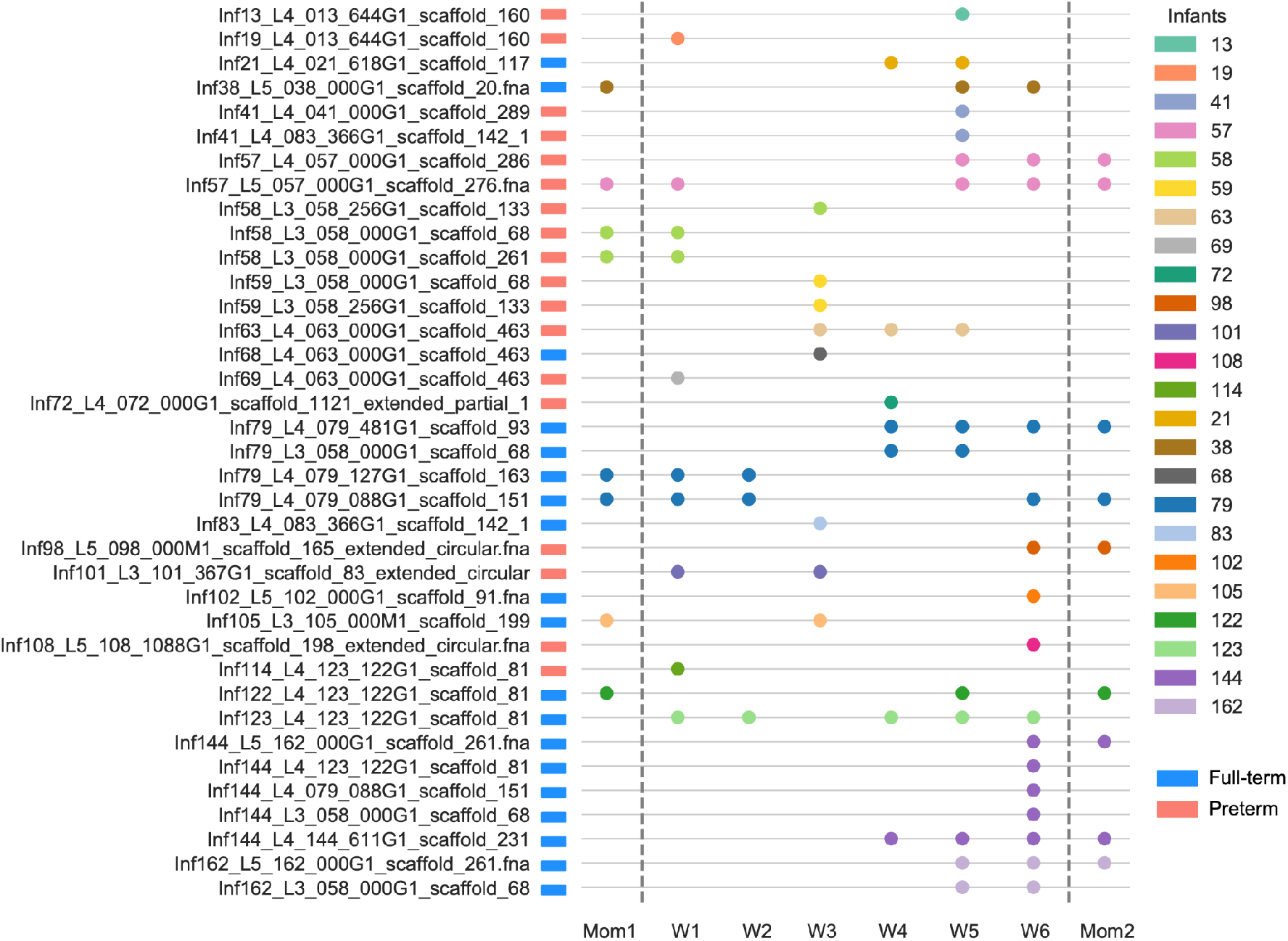
Colonization dynamics of recoded phages in infants and mothers. Schematic of all recoded phages in infants. Each row represents an infant-specific recoded phage strain, and circles represent the months in which the strain was detected. Strains are colored by infants. Circles in “Mom1” and “Mom2” indicate that the phages were vertically transmitted from the mother to the infant.

**Figure S8.**
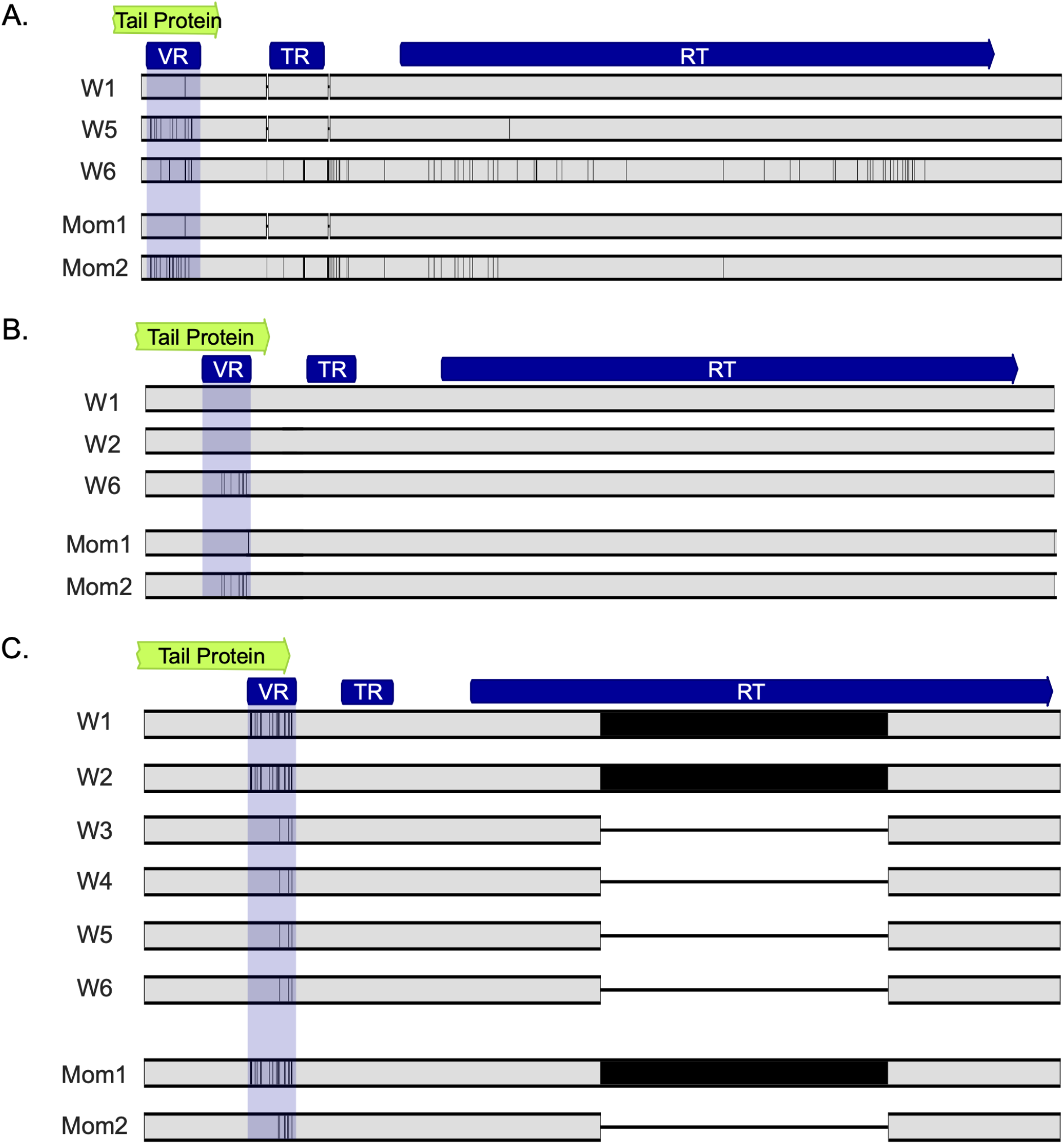
Nucleotide alignments of DGRs encoded by three recoded persisting phages. Sample-specific persisting phages of I57_PsAC1 (A), I79_PsAC1 (B), and I123_PsAC1 (C) were recovered, and their DGRs were aligned. Each row represents a sample-specific DGR region, and the solid vertical black lines represent SNPs that differ between sampling windows.

## Supplemental Tables

**Table S1. Infant metadata**

**Table S2. Details of metagenomics sequencing per fecal sample Table S3. Bacterial hosts enriched with phage persisters**

**Table S4. Bacterial genera enriched with bacterial persister strains**

**Table S5. Phage genes that accumulated a significantly high number of population-level mutation after 3-year persistence in infants**

**Table S6. Phage genes that accumulated a significantly high number of population-level mutation after 3-year persistence in mothers**

**Table S7. I57_PsAC1 genes with changed in-frame TAG frequency between the initial and final time windows**

**Table S8. Diversity-generating retroelements (DGRs) detected in three recoded persisting phages**

